# *Proteus*: an R package for downstream analysis of *MaxQuant* output

**DOI:** 10.1101/416511

**Authors:** Marek Gierlinski, Francesco Gastaldello, Chris Cole, Geoffrey J. Barton

## Abstract

*Proteus* is a package for downstream analysis of *MaxQuant* evidence data in the R environment. It provides tools for peptide and protein aggregation, quality checks, data exploration and visualisation. Interactive analysis is implemented in the *Shiny* framework, where individual peptides or protein may be examined in the context of a volcano plot. *Proteus* performs differential expression analysis with the well-established tool *limma*, which offers robust treatment of missing data, frequently encountered in label-free mass-spectrometry experiments. We demonstrate on real and simulated data that *limma* results in improved sensitivity over random imputation combined with a *t*-test as implemented in the popular package *Perseus*. Embedding *Proteus* in R provides access to a wide selection of statistical and graphical tools for further analysis and reproducibility by scripting. Availability and implementation: The open-source R package, including example data and tutorials, is available to install from GitHub (https://github.com/bartongroup/proteus).

## 1 Introduction

*MaxQuant* is one of the most popular tools for analysing mass spectrometry (MS) quantitative proteomics data (Cox and Mann, 2008). The output of a *MaxQuant* run usually consists of several tables, including the evidence data and summarized peptide and protein intensities. The downstream analysis and understanding of these data are essential for interpreting peptide and protein quantification. The standalone *Perseus* software package (Tyanova et al., 2016) is often used in conjunction with *MaxQuant* to help this interpretation.

The *Proteus* package described here offers simple but comprehensive downstream analysis of *MaxQuant* output in the R environment (R Core Team, 2018). The package is built with simplicity and flexibility of analysis in mind. A user unfamiliar with R can obtain differential expression results with a few lines code following the tutorial, while a more experienced R programmer can perform advanced analysis using the plethora of R and Bioconductor packages (Huber et al., 2015).

Differential expression is a commonly used term for statistical comparison of numerical results from two or more biological conditions. For high-throughput experiments, differential expression must take into account a statistical model of data distribution, wide range of variance, missing data and multiple test corrections. A number of tools have been developed to cope with these challenges, in particular in the field of RNA-seq (Gierliński et al., 2015; Schurch et al., 2016). One of these tools is the well-established Bioconductor package *limma* (Ritchie et al., 2015), originally written for microarrays (e.g. Peart et al., 2005), but often used with RNA-seq data (see Schurch et al., 2016, for comparison with other tools). The core feature of *limma* making it ideal for MS experiments is its ability to make analyses stable even for data with high proportion of missing values—this is achieved by borrowing information across features (that is transcript/genes in RNA-seq and peptides/proteins in MS). *Proteus* uses *limma* to perform stable and robust differential expression of data with gaps, thus avoiding the need for random imputation.

## 2 Data analysis in Proteus

*Proteus* analysis begins with reading the evidence file. To conserve memory only essential columns are retained. Reverse sequences and contaminants are rejected by default. In the current version, label-free, tandem mass tags (TMT) (Thompson et al., 2003) and stable isotope labelling by amino acids in cell culture (SILAC) (Ong et al., 2002) data are supported.

Peptide measurements (intensities or SILAC ratios) are aggregated from individual peptide entries with the same sequence or modified sequence. Quantification is carried out as the sum (label-free or TMT) or median (SILAC) of individual measurements. A user-defined function for peptide aggregation can be provided.

Protein intensities for label-free and TMT data are aggregated by default using the high-flyer method, where protein intensity is the mean of the three top-intensity peptides (Silva et al., 2006). For SILAC experiments, the median ratio is calculated. Alternatively, the sum of intensities or a user-provided function can be applied. The ability to aggregate peptide and protein data according to any prescription gives the package flexibility. On the other hand, the default, predefined aggregation functions make the package very easy to use. As an alternative to performing in-package aggregation, *MaxQuant’s* protein groups file can also be read directly into *Proteus*.

Peptide or protein data are encapsulated in an R object together with essential information about the experiment design, processing steps and summary statistics as mean, variance and number of good replicates per peptide/protein. Either object can be used for further processing, that is, analysis can be done on the peptide or protein level, using the same functions. Fig. 1 illustrates a few aspects of data analysis and visualisation in *Proteus*, using an example data set (see vignette/tutorial included in the package for details). These include peptide/protein count (Fig. 1A), sample comparison, correlation and clustering (Fig. 1B). Measurements can be normalized between samples using any arbitrary function, e.g., to the median or quantiles. A pair of conditions can be compared in a fold-change/intensity plot (Fig. 1C). The package provides functions to fetch protein annotations from UniProt servers.

**Figure 1:**
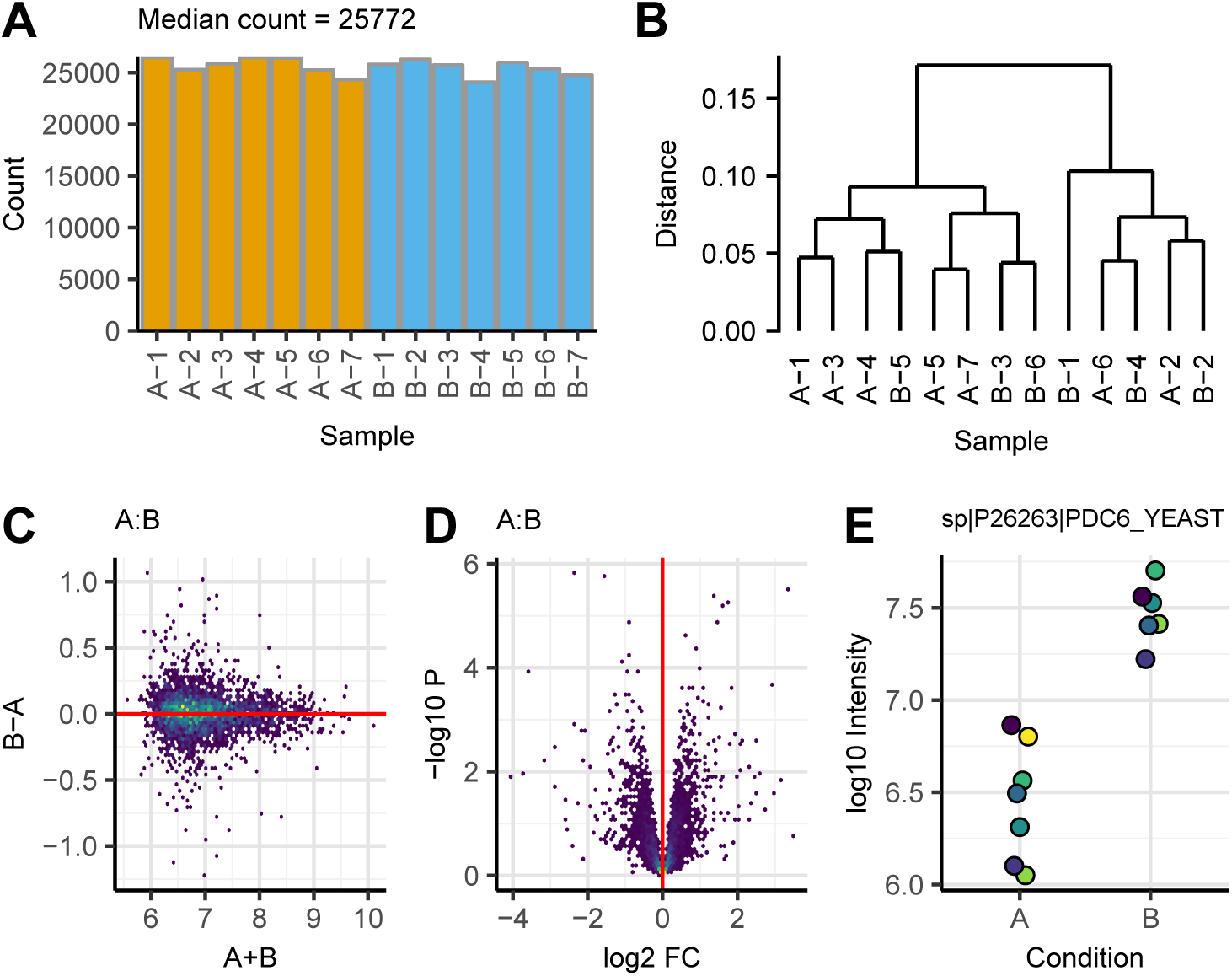
Visualization in Proteus using example data in two biological conditions (named A and B) and seven replicates each. This figure shows the actual plots created by Proteus. A. Peptide count per sample. B. Clustering of samples at protein level, a ’bad’ replicate B-7 was removed. C. Fold-change versus intensity for protein data. D. Volcano plot following differential expression analysis for protein data. E. Log-intensities of replicates (marked by different colours) in two conditions for a selected protein. The protein identifier, as extracted from evidence data, is shown at the top.

Package *limma* was chosen for differential expression due to its stability against missing data, common in label-free MS experiments (Lazar et al., 2016). *limma* offers an advantage over random imputation methods by borrowing information across peptides or proteins and using the mean-variance relationship to estimate variance where data are missing. The results can be visualised as a volcano plot (Fig. 1D) or as an intensity plot for individual peptide or protein (Fig. 1E).

*Proteus* offers a pointy-clicky data explorer based on the *Shiny* web application framework (Chang et al., 2018). It allows the properties of individual proteins to be studied in the context of the interactive volcano or fold-change-intensity plot (Fig. 2).

**Figure 2:**
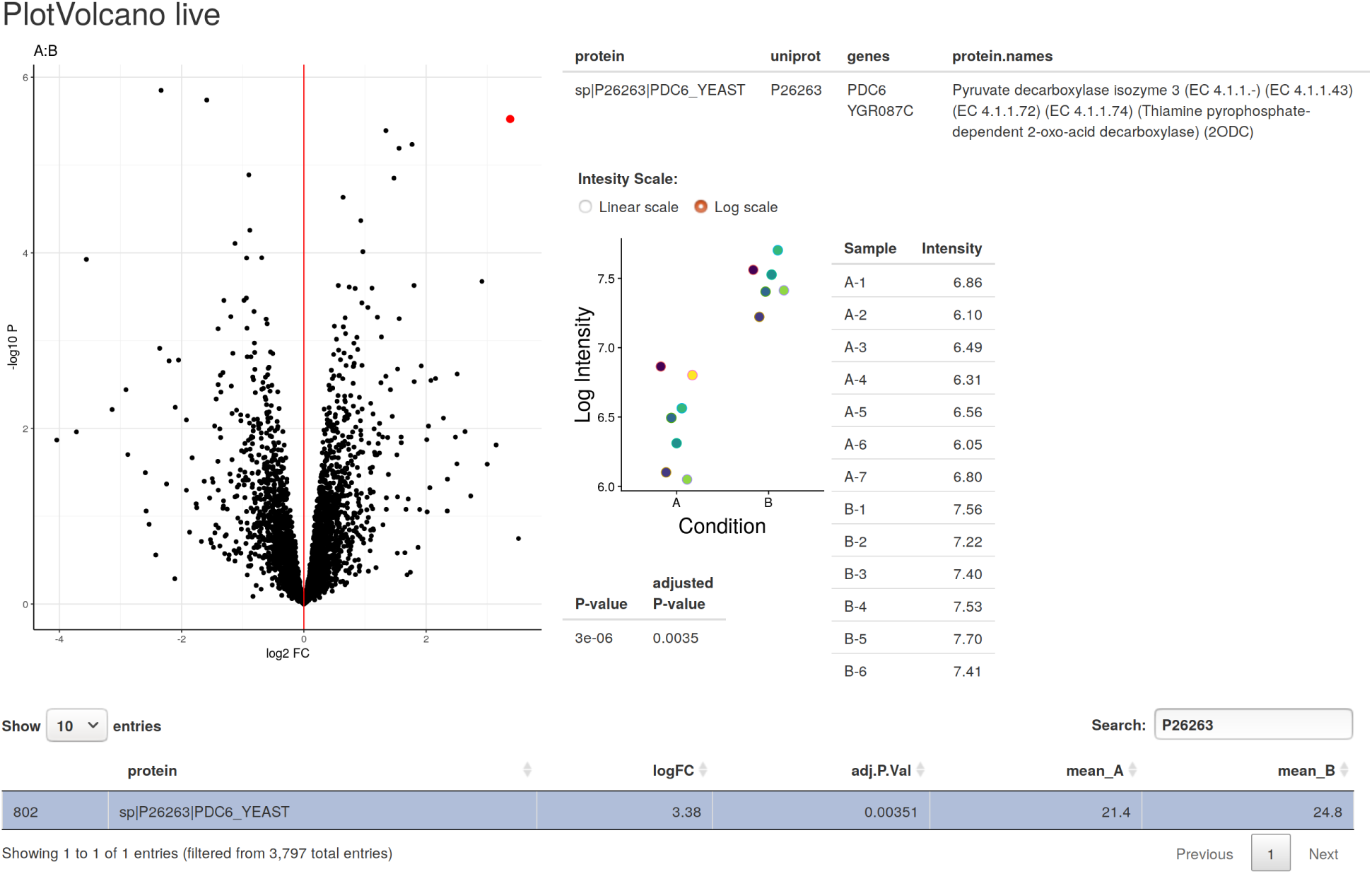
A screenshot of the interactive data explorer in Proteus, which is implemented in the Shiny framework. It shows an interactive volcano plot with a selected protein marked in red. The user can select proteins from the plot by hovering the mouse over a dot representing one protein. To the right, there are protein annotation, intensity plot, detailed intensity table and a p-value from the differential expression test. At the bottom there is a row selected from the full table of proteins by typing the protein ID in the search box.

### 2.1 Minimal example

A basic data processing flow in *Proteus* is as follows: read the evidence data and metadata, aggregate peptides and proteins, normalize protein data, perform differential expression and explore the results. Assuming that R variables `evidenceFile` and `metadataFile` point to *MaxQuant’s* evidence file and a small text file describing the design of the experiment (that is how samples and conditions relate to evidence data) the minimal R code to process these data is:

~~~
# read evidence data
evidence <-readEvidenceFile(evidenceFile)

# read metadata
metadata <-read.delim(metadataFile, header=TRUE, sep=“\t”)

# aggregate peptides
peptides <-makePeptideTable(evidence, metadata)

# aggregate proteins
proteins <-makeProteinTable(peptides)

# normalize protein intensities
proteins <-normalizeData(proteins)

# differential expression
res <-limmaDE(proteins)

# interactive data expolorer
plotVolcano_live(proteins, res)
~~~

Here default parameters were applied in all steps, but each function has several parameters allowing full control over, e.g., the way peptides and proteins are aggregated or normalized. *Proteus* comes with a set of tutorials (R vignettes) using real data examples to illustrate every step of data processing. Each function in the package is accompanied with detailed documentation and examples.

## 3 Proteus vs Perseus

Here we compare the performance of *Proteus* (version 0.2.8) to *Perseus* (version 1.6.1.3), a commonly used *MaxQuant* data analysis tool with a graphical user interface, available for MS Windows, on two examples. First, we analyse a label-free proteomics data set in two conditions and three replicates each. Second, we create a simulated data set based on statistics from a large real data set, to investigate power and false positives from both tools. We focus on the performance for detection of differential expression, which is carried out by a t-test in *Perseus* and *limma* in *Proteus*.

### 3.1 Simple data set

First we compared differential expression offered by both packages using the same protein data set. We used a subset of a large data set from Gierlinski et al. (in preparation). We read the evidence file and filtered out reverse sequences and contaminants. Then, we created a peptide table based on three randomly selected replicates in each condition (samples Mut-7, Mut-28, Mut-34, WT-8, WT-13, WT-25). Peptides were aggregated by summing multiple evidence entries for a given (unmodified) sequence. Next, we aggregated peptides into proteins using the high-flyer method and peptide-to-protein mapping based on the ’leading razor protein’ column from evidence data. We filtered protein intensities in these replicates, so at least one data point was present in each condition. These data were normalized to median (that is, after normalization median intensity in each sample was the same). The resulting intensity table containing 3338 proteins in two conditions in three replicates each was processed in *Perseus* and *Proteus*.

In *Perseus* we log_2_-transformed these data and filled missing values with random imputation, using the default parameters, width = 0.3 and down shift = 1.8. Then, we performed a two-sample *t*-test and exported the results as a generic table. In *Proteus* we log_2_-transformed data and performed differential expression using *limma*. Since the protein intensities were the same, we compared the difference between *t*-test with imputation versus *limma* without imputation.

Figure 3 shows the comparison of p-values, significantly differentially expressed proteins and volcano plots for *Perseus* and *Proteus*. There were 39 proteins called as significant by both methods, 15 only by *Perseus* and 45 only by *limma* (in *Proteus*), see Fig. 3B. We can see in Figure 3C a small group of proteins called by *limma* only where *Perseus* reported large p-values (6 blue circles at the bottom of the plot marked as “1”). These proteins have missing data and rather large intensity. Imputation in *Perseus* filled the missing values with low intensities, inflating variance and missing what otherwise would be differentially expressed. An example of such a protein is shown in Figure 4A. The other group of *limma*-only blue circles in Figure 3C (marked as 2) indicates that the permutation FDR method used in *Perseus* is slightly more conservative that that in *limma*. All these proteins have adjusted p-values near the limit 0.05. An example is shown in Figure 4B. The proteins plotted as green triangles (see Figure 3 C and D) are marked as differentially expressed by *Perseus* but not *limma*. They typically have small variance and small fold change. They are called significant by a simple t-test but not by *limma*, which moderates variance and avoids cases of unusually small variability. An example of such a protein is shown in Figure 4C. We note that data with small fold changes can be easily removed from *Perseus* by setting a fold-change limit in the *t*-test.

**Figure 3:**
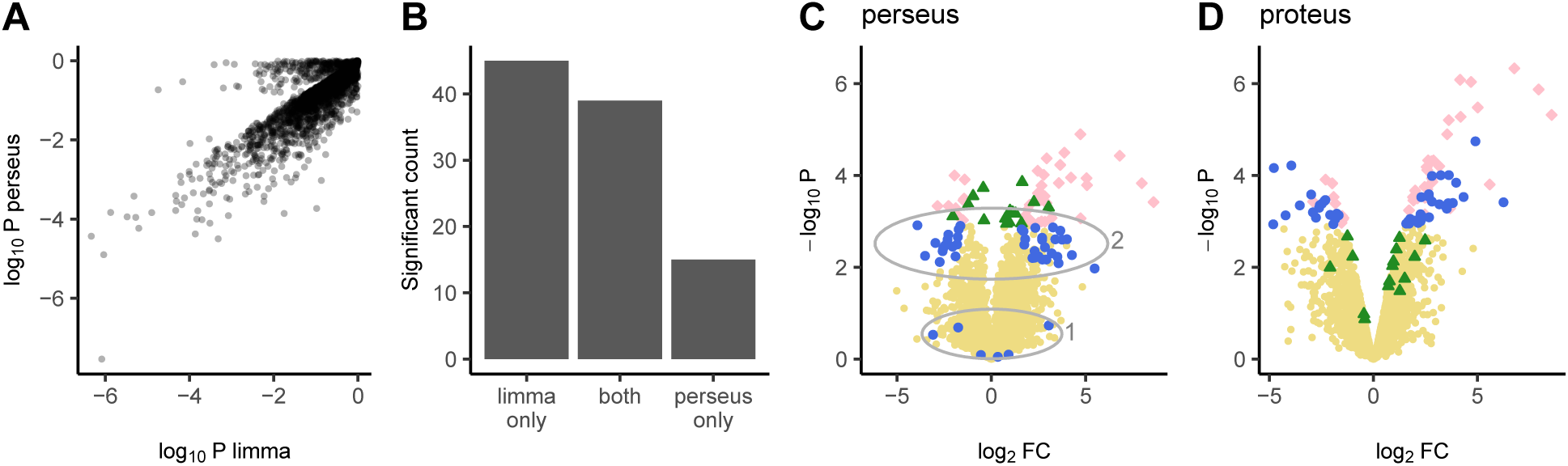
Perseus vs limma for a selection of 3 vs 3 replicates. A. P-value (not adjusted) comparison. B. Number of significant proteins. C. Volcano plot using fold-change and p-values from Perseus. All data are in yellow background. The limma-significant-only proteins are as blue circles, the perseus-significant-only proteins are as green triangles. Proteins significant in both tools are as pink diamonds. The ellipses indicate data discussed in the text. D. Volcano plot using fold-change and p-values from Proteus. Symbols are the same as in C.

**Figure 4:**
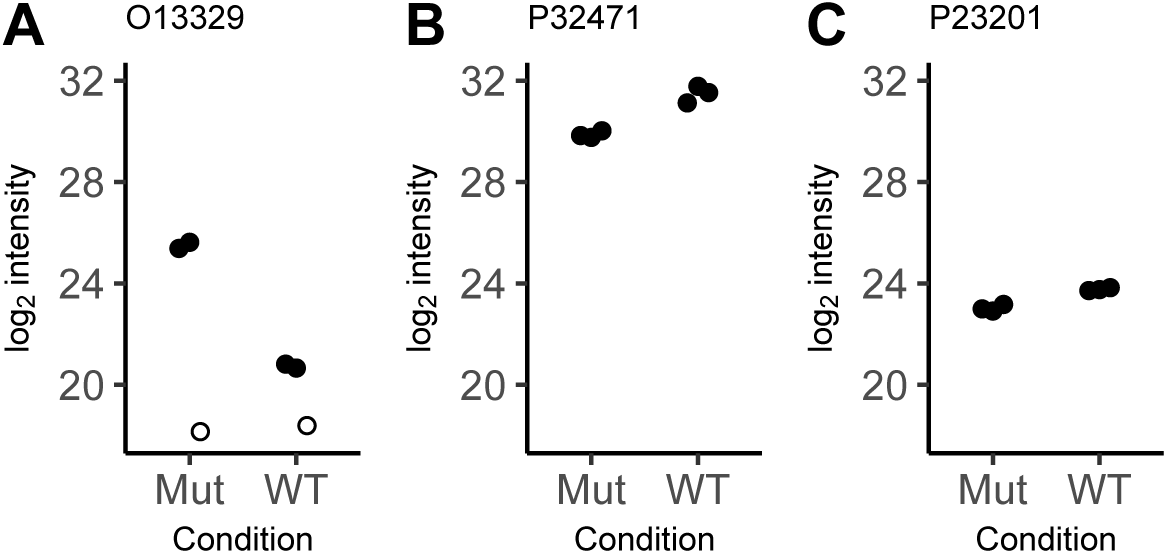
Selected examples of proteins called as significant by one tool only. UniProt identifiers are shown on top of each panel. A. Called by limma only. Imputation in Perseus inflated variance creating a false negative. B. Called by limma only. Perseus FDR is more conservative than that in limma at the same limit of 0.05. C. Called by Perseus only. An example of very low variance, which is moderated and called negative by limma. Data imputed by Perseus are marked with open circles.

The imputation in *Perseus* is designed to fill missing low-intensity data with a randomly generated Gaussian numbers (see supplemental figure 3 in Tyanova et al. (2016)). However, on some occasions a datum can be missing even at high intensities. In such cases variance is dramatically inflated and the protein is not called as differentially expressed. We warn against using data imputation. *limma* offers a better approach to missing data, by modelling mean-intensity variance and using moderated variance for the test. Certainly, the imputation step can be omitted in *Perseus*, but this reduces power and rejects data with only one replicate available in a condition. Again, *limma* can estimate variance and make a decision about differential expression even in such extreme cases (at an increased risk of a false positive).

### 3.2 Simulated data

We next compared performance of differential expression in both tools using simulated data. We generated a simulated set based on real data. Since we have a good data set of two conditions in 35 replicates each (Gierlinski at al. in preparation), we used it to find the mean-variance relationship and the rate of missing values as a function of the intensity. We used this information to create a simulated data set in two conditions in three replicates each and allowed for missing data in each condition. We chose a grid of log_2_ *M* (mean) and log_2_ *FC* (fold change) covering most of the original data. For each combination of log_2_ *M* and log_2_ *FC* we generated two random samples of up to 3 data points from the log-normal distribution with the given mean and variance estimated from the linear function found from real data. The first sample had the mean *M*, the second sample had the mean *M* * *FC*. For each sample the number of good replicates was generated to mimic the real data. First, for the given *M*, we used the cumulative distribution of the number of good replicates to generate a number between 1 and 35. This was then sub-sampled to the 3 replicates generated (for example, if 10 was generated in the first step, we created a vector of 10 good and 25 bad replicates and drew a random sample of 3). Since we are not interested in samples with no data, we enforced at least one good replicate in each sample. This means that data with only one good replicate will be over-represented for very low intensities. This is not an issue as our aim is to assess tool performance at each intensity level and low intensities will invariably contain a lot of missing data. For each combination of log_2_ *M* and log_2_ *FC* we generated 1000 samples in two conditions, using this technique. This gave us a large set of 105,500 “proteins” covering a wide range of intensities and fold changes.

We performed differential expression on the simulated data using *Perseus* and *Proteus*. In *Perseus* we imported simulated data from a file, log_2_-transformed, applied default imputation and used a two-sample *t*-test. In *Proteus* we log_2_-transformed the data and used *limma* for differential expression. The results are shown in Figure 5. Panels A and C show the proportion of proteins called significant in a group of 1000 proteins for each combination of fold change and mean intensity. We can see that *limma* (in *Proteus*) performs well across all intensities, discovering almost all positives for the highest log_2_ *FC* = 2.8 used here. In contrast, the sensitivity of *Perseus* drops dramatically at low intensities. Even at medium intensities of log_2_ *M* = 23 only about half of the changing proteins are discovered at large fold changes of log_2_ *FC* = 2. See also Figure 7A.

**Figure 5:**
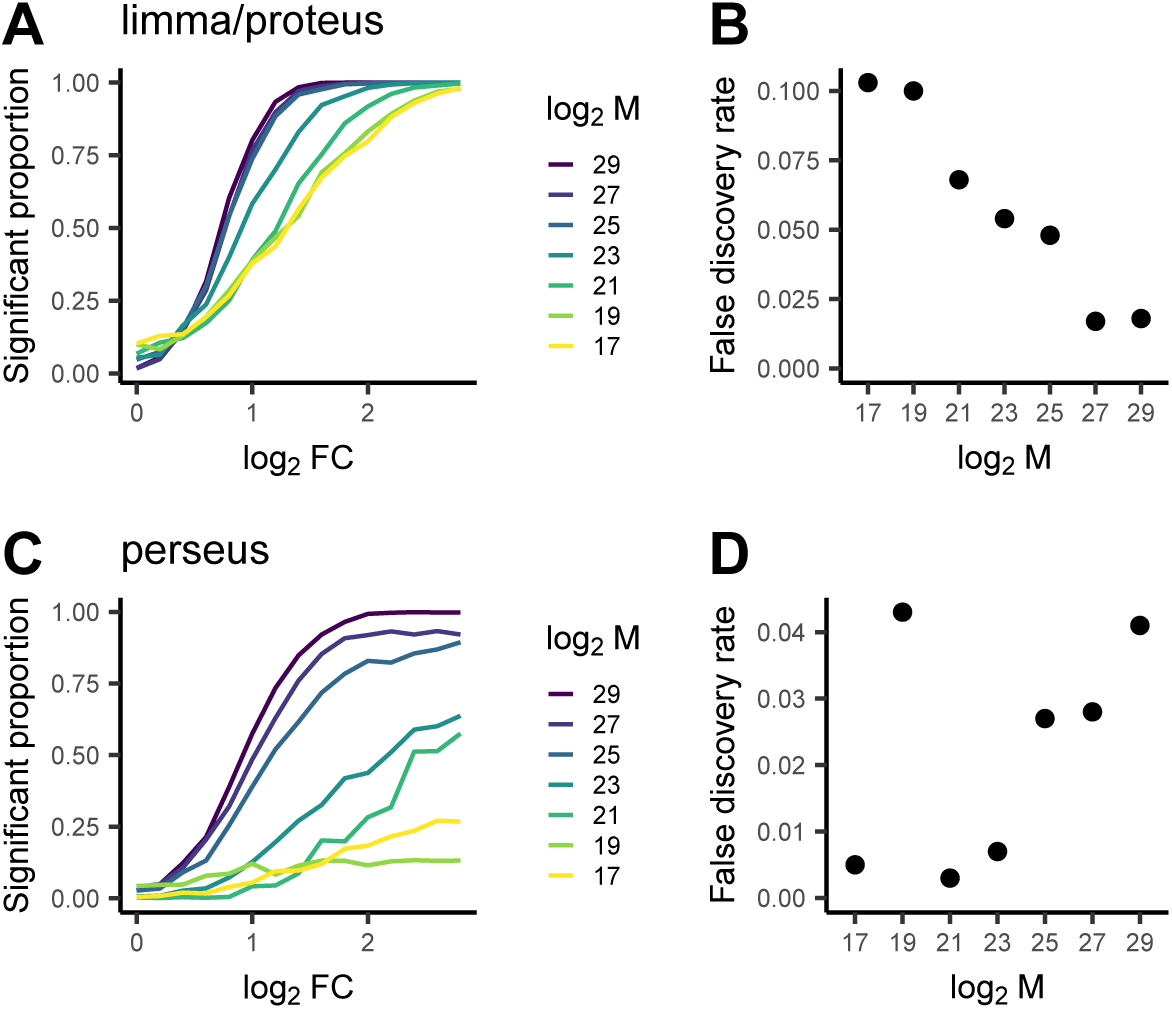
Results for the full set of simulated data with 3 replicates. Top panels show limma, bottom panels show Perseus results. A and C show the proportion of tests called as signficant as a function of the simulated fold change (FC) and mean (M). B and D show the false discovery rate, that is the proportion of tests for simulated log FC = 0 called as significant.

**Figure 7:**
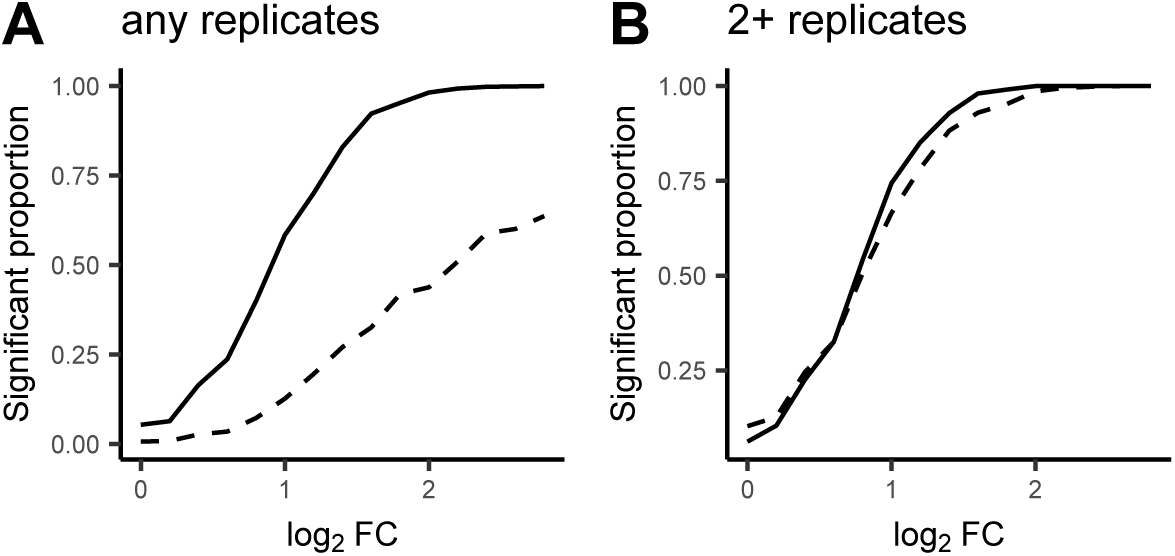
Comparison of significance curves corresponding to log M = 23. A. Full simulatred data set (see Figs. 5A and 5C). B. Filtered simulated set (see Figs. 6A and 6C). Perseus results are shown in dahsed curves, Proteus results are represented by solid curves.

The main reason for this behaviour is imputation of missing replicates in *Perseus*. We notice that due to the way simulated data were generated, all proteins for the lowest intensity log_2_ *M* = 17 contain only one good replicate in each condition. As the *t*-test cannot deal with samples of one, imputation is necessary and the result is randomized. On the other hand, *limma* borrows information across the entire set and builds a reliable model of variance which works for any sample size. As we can see from the bottom curve in Figure 5A (corresponding to log_2_ *M* = 17) *limma* performs well even in tests of one versus one replicate.

The increased sensitivity of *limma* results in an increased false discovery rate (FDR). We can estimate FDR as a proportion of proteins called significant at log_2_ *FC* = 0. Figure 5B shows that FDR for *limma* exceeds the assumed limit of 0.05 at the three lowest intensities. *Perseus* discovers far fewer positives at these intensities, which results in lower FDR (Figure 5D).

Since imputation is clearly an issue we decided to compare *Proteus* and *Perseus* using data that do not require imputation. We used the same simulated data set, but filtered out all proteins with only one good replicate in either condition. Filtering low-replicate data would reflect a more realistic workflow for a researcher who doesn’t want to apply imputation. After filtering, we processed data in *Perseus* and *Proteus* as before, but skipped the imputation step in *Perseus*. Results are shown in Figure 6. Not surprisingly, the significant proportion of *limma* and *Perseus* are now more similar, though *limma* still offers slight advantage (see also Figure 7B). The false discovery rate is now better controlled by *limma* than by *Perseus* where 4 out of 5 intensity groups result in *FDR* ∼ 0.1.

**Figure 6:**
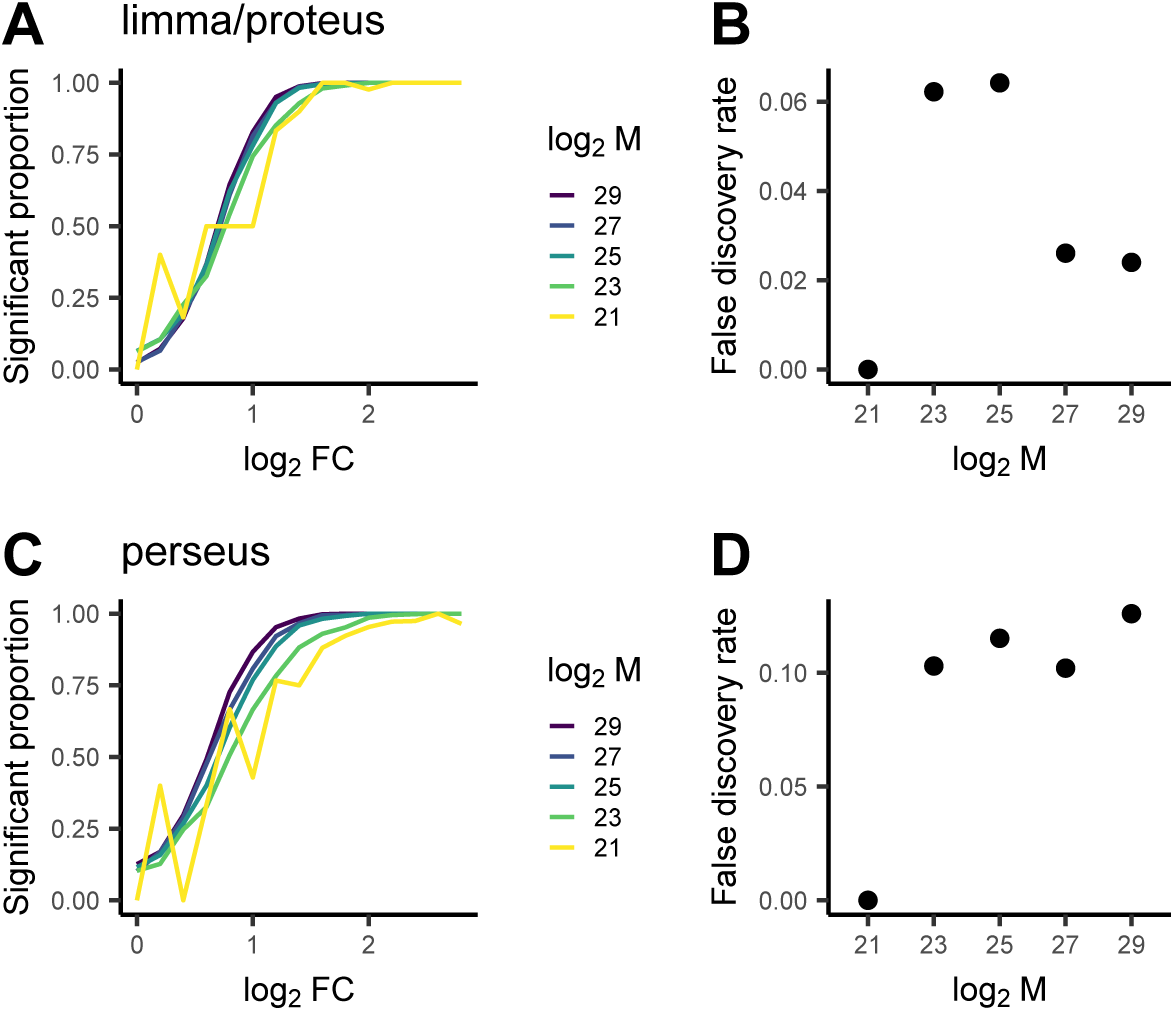
Results for the filtered set of simulated data with 3 replicates. Only data with at least 2 replicates in each condition were used. Panels are the same as in Figure 5.

## 4 Conclusions

R is becoming one of the most widely used tools for data science and statistical computing, in particular in academia (Tippmann, 2014; Muenchen, 2017). Its strength is built on the wealth of statistical libraries available. *Proteus* adds to a rapidly growing suite of bioinformatics packages in R. It not only performs specific tasks related to processing of *MaxQuant* output, but opens peptide and protein data to further analysis and visualisation. It offers an alternative to the popular package *Perseus* for researchers wishing to explore the benefits of the R environment.

R is a scripting language and when data and code are published together, it makes data processing fully reproducible. Any analysis performed in *Proteus* can be replicated by any researcher, including all intermediate steps, simply by running the original code again. We recommend using the *RStudio* environment (RStudio Team, 2015), where the code can be executed step-by-step and each R object can be easily scrutinised. R is a cross-platform project and can be used on most operating systems. Needless to say, *Proteus* is fully open source.

*Proteus* uses a powerful package *limma* for differential expression analysis, allowing for stable analysis of data with missing values, common in label-free MS proteomics, with no need for random imputation. Instead, *limma* borrows information between peptides/proteins to build a robust model of variance and performs differential expression tests based on this model. We demonstrate that this offers a clear advantage in terms of sensitivity over a *t*-test combined with random imputation. This makes *Proteus* particularly useful for label-free data with a small number of replicates.

## 5 Acknowledgements

The authors would like to thank Katharina Trunk, Sarah Coulthurst and Julien Peltier for kindly providing example data used in the *Proteus* package and discussed in this paper. MG thanks Matthias Trost for discussions and James Abbott for support.

